# Predicting host associations of the invasive spotted lanternfly on trees across the USA

**DOI:** 10.1101/2022.09.12.507604

**Authors:** Nicholas A. Huron, Matthew R. Helmus

## Abstract

Global impacts of invasive insect pests cost billions of dollars annually, but the impact of any individual pest species depends on the strength of associations with economically important plant hosts. Estimating host associations for a pest requires surveillance field surveys that observe pest association on plant species within an invaded area. However, field surveys often miss rare hosts and cannot observe associations with plants found outside the invaded range. Associations for these plants instead are estimated with experimental assays such as controlled feeding trials, which are time consuming and for which few candidate hosts can be tested logistically. For emerging generalist pests, these methods are unable to rapidly produce estimates for the hundreds of potential suitable hosts that the pest will encounter as it spreads within newly invaded regions. In such cases, association data from these existing methods can be statistically leveraged to impute unknown associations. Here we use phylogenetic imputation to estimate potential host associations in an emergent generalist forest pest in the U.S., the spotted lanternfly (*Lycorma delicatula*; SLF). Phylogenetic imputation works when closely related plants have similar association strengths, termed phylogenetic signal in host association, which is common in phytophagous insects. We first aggregated known SLF host associations from published studies. Existing research has estimated association strengths for 144 species across both the invaded and native range of SLF. These known associations exhibited phylogenetic signal. We then developed two protocols that combined known host association data and fit phylogenetic imputation models based on hidden state prediction algorithms to estimate association strength for 569 candidate tree species found across the continental U.S. Of candidate species considered, 255 are predicted to have strong associations with SLF in the U.S. and can be found in several clades including Juglandaceae, Rutaceae, Salicaceae, and Sapindaceae. Uninvaded regions with the highest numbers of these strongly associated species include midwestern and west coast states such as Illinois and California. Survey efforts for SLF should be focused on these regions and predicted species, which should also be prioritized in experimental assays. Phylogenetic imputation scales up existing host association data, and the protocols we present here can be readily adapted to inform surveillance and management efforts for other invasive generalist plant pests.

## Introduction

As the Earth continues to become more interconnected and interdependent through human activities, invasive detrimental insects (insect pests) will continue to grow as impediments to agricultural and forestry production globally^1–7^. Insect pests may pose a great threat to global biodiversity and ecosystem functioning, because they are spread easily via human trade networks^3,8,9^ and their introductions have considerable impacts. Insect pests of plants stand to induce crop loss, thereby producing shortfalls in food and medicine supply, fiber and building material stores, and biofuel reserves^6,10^. Such impacts are estimated to cost over 70 billion USD per year globally, with nearly 10% of cost localized to the U.S.^11,12^.

Impacts depend on the breadth of host plants that a pest associates with (e.g., via feeding, sheltering, and egg laying) and the strength of such associations^13–15^. Insect pests can range from specialists with few host species, such as the emerald ash borer (*Agrilus planipennis*), which feeds on ash trees (*Fraxinus* spp.) and is responsible for the loss of over 25 millions of acres of forest in the U.S. between 2014 and 2018^16,17^, to the spongy moth (*Lymantria dispar*), a generalist with over 500 host plants and is responsible for the loss of over 6 million trees over the same five-year span^17–19^. Generalists associate with many different host species, but the comparative strength of association among host species can vary greatly and management must account for this variability^20^. Often this variability is unknown when a pest is introduced to a region with suitable host species that it has not encountered before.

Pest associations are estimated using two different methods: field surveys and experimental assays^13,20^. Observations from field surveys identify host species, confirm associations for a new introduction, and justify initial management plans^20^. By contrast, experimental assays validate observations made in the field and investigate associations with species not observed in the field more closely^13^. In doing so, experimental assays can identify the comparative strength of relationships between a pest and members of its host species guild and ultimately uncover the factors that drive association preferences^21^. Although both approaches are vital to estimate association strength and inform management, using only these two methods has drawbacks. First, field surveys are biased because they sample locations to determine where a pest is or is not, thereby sampling abundant plant host species more than rare ones. Furthermore, they must balance standardized methodology and optimal timeframes for sampling, which either limits the questions they can address or can produce inconsistent results. Second, experimental assays are resource intensive and can fail to assess important species without appropriate background knowledge, meaning that it can take a lot of time before they can accurately contribute to management plans^13,21^. Given these drawbacks to both methods, it is unsurprising that even though databases are improving (e.g., ^21,22^), plant host associations for insect pests remain poorly documented^23,24^. Unlike species constrained to their native range, introduced pests can have distributions not at equilibrium, encounter novel candidate host species, and experience ecological release^25–28^. These details are important for pest management but are often poorly understood, especially early in an invasion. Therefore, it is necessary to seek additional methods for studying insect pest host associations, especially for emergent invasions.

Imputation is a statistical process to estimate unknown data from observed relationships among known data and other covariates. Phylogenetic imputation methods predict species traits based on shared evolutionary history^23,29–31^. Numerous phylogenetic imputation approaches exist^32,33^, all of which assume the trait of interest demonstrates phylogenetic signal, where closely related species have similar traits^34–36^. To date, phylogenetic imputation has been used to predict biochemical, ecological, morphological, and physiological traits for contemporary and ancestral species^36,37^. Host associations for insect pests often exhibit phylogenetic signal^21,38–41^, which can be considered a proxy for latent host variables that demonstrate phylogenetic signal as well^42,43^, such as morphology, phytochemicals, and life history^21,44,45^. Phylogenetic signal has been suggested as a relationship to leverage for identifying novel host associations^21,46,47^, but no empirical work to date has used phylogenetic imputation to explicitly estimate potential host associations to inform monitoring and management for a high-risk pest.

The spotted lanternfly (SLF, *Lycorma delicatula*, Hemiptera, Fulgoridae) is a phytophagous forest pest planthopper that has been introduced outside its native range at least three times to date: South Korea and Japan in the early 2000s and then to Pennsylvania in the U.S. c. 2014^48^. Since its U.S. introduction, SLF has spread rapidly throughout the eastern U.S. and continues to spread to other states^49,50^. Continued rapid spread suggests that SLF is likely to have further introductions^49^, including via stepping-stone invasions to new regions^2,51^. Part of why SLF has spread so quickly in the U.S. may be because it is a generalist with a wide range of hosts^51^. SLF is known to feed on over 100 plant species worldwide, including economically important trees such as apples, cherries, maples, and walnuts^48,52–54^. When heavy SLF feeding takes place, the tree can flag, mold can accumulate and branches can die back, all of which contribute to economic damage^48^.

SLF associates more readily with some hosts over others^51^. Several preferred hosts have been identified, such as tree of heaven (TOH, *Ailanthus altissima*)^53^, but not all hosts appear to support SLF throughout its lifecycle^55^. In fact, it was initially unclear if SLF needed to feed on TOH for part of its life cycle to complete development^56^, but more recent research indicates that SLF can develop on several other woody tree hosts (e.g., black walnut [*Juglans nigra*], butternut [*J. cinerea*], chinaberry [*Malia azedarach*], Chinese toon [*Toona sinensis*], and tulip tree [*Liriodendron tulipifera*]) and non woody hosts (e.g., grapes [*Vitis labrusca, V. vinifera*], hops [*Humulus lupulus*], and Virginia creeper [*Parthenocissus quinquefolia*])^51,56,57^. SLF preference for this diverse array of known hosts could be due to variation in phytochemicals, plant sugar content, or other phylogenetically conserved traits such as venation structure^42,43,53,58–60^, but it is unclear which of the above contribute to variation in SLF host association or the extent to which SLF prefers one host over another^51,56^.

As SLF spreads, additional hosts that it will readily associate with for some or all its life cycle will be discovered. Therefore, it is prudent to elucidate patterns of known and potential SLF host association, especially for trees, because SLF has already demonstrated a preference for several woody host species that are important food crops, sources of lumber, or cultivated ornamentals. If additional suitable hosts are not identified promptly, there may be consequences to affected industries. Concern over SLF is clear, as considerable resources have been allocated by U.S. state and federal governments^51,61,62^ to better understand this pest and its host plants. As such, host association data measured by traditional methods (e.g., field surveys and experimental assays) are increasing but likely to remain incomplete for some time.

Here, we present a phylogenetic imputation approach to ask which novel tree species are predicted to be strongly associated with SLF as it spreads throughout the U.S. (Figure 1), given the high proportion of known hosts that are trees. We then set high risk trees within an applied management context by building a map of the U.S. to identify areas with many known and likely host species for SLF that should be monitored.

**Figure 1:**
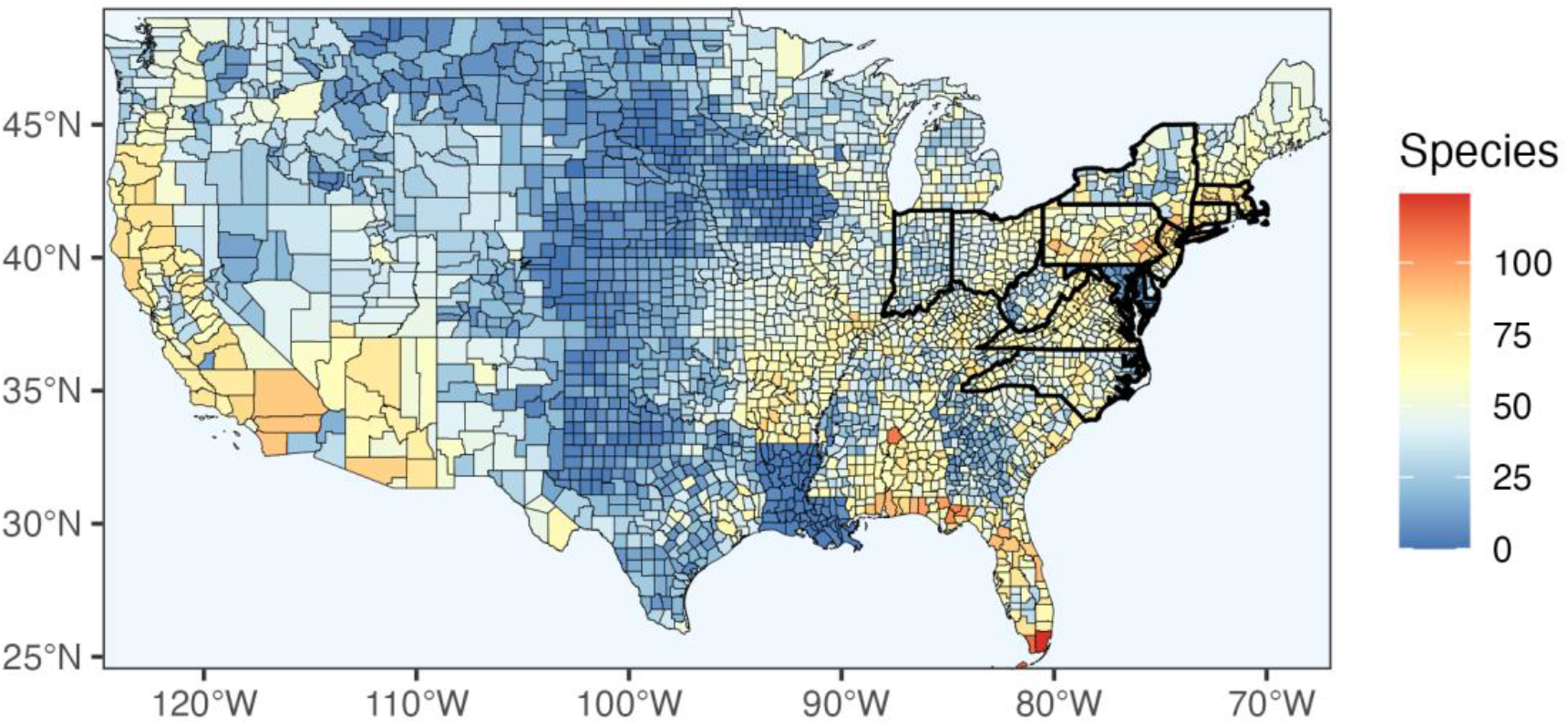
Map of candidate tree species by county. Spotted lanternfly (*Lycorma delicatula*; SLF) candidate host tree species are distributed throughout the contiguous U.S. Candidate species richness is concentrated heavily in established regions for SLF (mid-atlantic and northeast, states outlined boldly) but also includes uninvaded regions.

## Methods

### Known Host Association Strengths and Candidate Hosts

We searched the literature for studies on SLF host association by querying Google Scholar (https://scholar.google.com) and Web of Science (https://www.webofscience.com) for *Lycorma delicatula* or spotted lanternfly as well as reviewing SLF-specific collection of studies produced by the SLF Working Group (https://stopslf.org) to identify those that compared associations across species. We sought studies in the following categories: field survey observations, survival experimental assay, and decision experimental assay, which resulted in 16 studies^54–56,60,63–73^. We then screened studies to find those that compared numerous species and looked at associations for individual hosts rather than combinations. We contacted authors for data when the data in a relevant study were not public. Wherever possible, we used studies that include comparisons of a variety of plant species and across SLF life stages to maximize knowledge of host associations. After screening, five final host association studies remained for our analyses.

We combined unique species across all retained studies to determine a list of known SLF host species. To use all available data for measuring association, we included all known woody and non-woody hosts. Species names were screened for taxonomic updates with the R package taxize^74^ v. 0.9.99 to ensure proper species and family-level relationships among them (and candidate hosts later) according to the Integrated Taxonomic Information System (ITIS)^75^. Some known host taxa were still represented ambiguously for some studies (e.g., *Quercus* spp. in Dechaine et al.^70^), because the authors did not specify which species within the genus was used. We addressed these taxa by identifying known hosts that are congeners present in other studies and geographically plausible (recoding: *Arctium* spp. = *Arctium lappa, Fraxinus* spp. = *Fraxinus americana, Quercus* spp. = *Quercus rubra, Sassafras* spp. = *Sassafras albidum, Syringa oblata* var. *dilatata* = *Syringa oblata*, and *Vitis* spp. = *Vitis vinifera*).

We then obtained observed measures of SLF host association from the retained studies. For those with continuous data (Table 1), we calculated the mean association measures for each plant species that spanned as many life stages or length of season as possible. If data were reported for multiple seasons (e.g.,^70^), we averaged measured associations across all reported seasons. The decision to group life stages or seasonal sampling together was motivated by a desire to standardize association data and provide an initial assessment that can be refined as finer resolution data become available. In practice, association was most often a measure of the proportion of individuals that chose a particular host over all others assessed (Table 1). We also considered one other continuous association measure, mean longevity per host species^60^. Available studies that measured continuous association were unable to incorporate many known hosts, and so we also created a dataset derived from the list of known host species reported in Barringer and Ciafré^54^. Their results include broad SLF life stage use of particular hosts (egg, nymph, and adult), which we converted into an ordinal measure of association by counting the number of life stages that use a particular host, thereby rounding out our SLF host association dataset (Table 1).

**Table 1:** Summary of known host studies data. Individual studies used to predict host association from known plant host species for spotted lanternfly (*Lycorma delicatula*; SLF). The number of hosts with measured SLF association included many tree species and differed across studies. Specific study citations can be found in the main text references.

| Study Name | Hosts | Measured Hosts | Measured Tree Hosts | Measure | Study Type | Life Stage Coverage |
| --- | --- | --- | --- | --- | --- | --- |
| Barringer and Ciafré 2020 (barringer1) | 157 | 127 | 112 | ordinal number of life stages (1–3) | field survey | all |
| Dechaine et al. 2021 (dechaine23) | 31 | 31 | 26 | continuous log total relative proportion | field survey | all |
| Derstine et al. 2020 (derstine_sum) | 14 | 14 | 10 | continuous proportion of yes:no choices (negative control) | decision experiment | incomplete all |
| Kim et al. 2011 (kim_sum) | 13 | 13 | 13 | continuous log total relative proportion | field survey | 3rd–adult |
| Murman et al. 2020 (murman4) | 25 | 25 | 23 | continuous mean longevity per host | survival experiment | all |
| Total Unique | 174 | 144 | 125 |  |  |  |

Since SLF is known to feed on numerous tree species, including those that are economically important, we focused our efforts here on candidate host tree species found within the U.S. To obtain a list of candidate host species, we began with an inventory of tree species found in the U.S.^76^ This species list combines several published surveys of tree species^77–81^, which were aggregated but preserved in their original form. We then updated family and species-level taxonomy as for known species. Given that this dataset did not include some agriculturally important tree species (e.g., avocados, *Persea americana*), we also queried the USDA National Agricultural Statistics Service (USDA NASS) Quickstats database^82^ on 28 June 2022 for additional tree species that are cultivated domestically to add as candidate hosts for consideration.

### Host Plant Phylogeny

To build an appropriate phylogeny of all known and candidate SLF hosts, we harnessed a recently published R package, V.PhyloMaker^83^ v. 0.1.0. V.PhyloMaker provides convenient means to prune or graft tree branches based on the most recently updated mega-trees that show relationships among extant seed plants. We combined known and candidate host species lists and extracted a phylogeny using the default backbone megatree, GBOTB.extended.tre^83^, which updates taxonomy for and combines a recent seed plant phylogeny^84^ with one for pteridophytes^85^. There were 176 species not in the phylogeny, which we grafted on the tree under V.PhyloMaker scenario 3, an author-recommended approach for robust species-level phylogenetic analyses^86^ that binds new families or genera at branch distances based on relative clade branch lengths and new species to the basal node for its genus (see ^83^ for details).

Preliminary phylogeny construction was checked to identify potential taxonomic inconsistencies between the backbone megatree and our species lists. As a result, we revised the family placement of several genera (changed to Family: Genus as follows: Cucurbitaceae: *Luffa*, Malvaceae: *Firmiana*, Malvaceae: *Tilia*, Moraceae: *Broussonetia*, and Nyssaceae: *Nyssa*). We also treated several synonyms in the backbone tree as species in the known and candidate host dataset without more explicit correct tips, which were altered for phylogeny construction but returned to their original state for future analyses (host dataset = backbone phylogeny as follows: *Salix koreensis* = *S. pierotii, Chamerion angustifolium* = *Epilobium angustifolium, Humulus japonicus* = *H. scandens, Platanus x acerifolia* = *P. hispanica, Prunus x yedoensis* = *Cerasus yedoensis, Pseudocydonia sinensis* = *Chaenomeles sinensis, Rosa hybrida* = *Rosa* hybrid cultivar). By making the above changes, we reduced the number of ad hoc grafts to our phylogeny while likely placing such species in the tree most appropriately, given current taxonomic understanding.

### Phylogenetic Imputation Protocols for Predicting Association

We developed two protocols for combining known host association data across multiple studies and using phylogenetic imputation to predict candidate host association strengths. Hidden State Prediction (HSP)^36^ is a phylogenetic imputation method related to ancestral state reconstruction and phylogenetic comparative methods. We chose HSP over other methods because it relies on the entire phylogenetic topology to predict terminal tips.

Our first HSP protocol (HSP1) estimated host associations with data from each study individually and then combined the results as a composite association metric. This protocol allowed us to use both continuous and ordinal data, because we fit a separate model for each of the data sets that came from the association studies. For HSP1 continuous data, we added the lowest value for each study and log10 transformed the data. The states for each study were rescaled and then combined. Our second HSP protocol (HSP2) averaged the continuous data studies and then conducted HSP with the average known association data. The one ordinal data study, Barringer and Ciafré 2020^54^, was not included in this protocol. Continuous data for HSP2 were similarly log10 transformed and rescaled (see below) and then averaged for each species across studies with known host data before their use in HSP.

We used the R package castor^33^ v. 1.7.2, which provides HSP methods for both continuous and ordinal data. For continuous association data, we performed HSP with phylogenetic independent contrasts^29^ under default settings (castor::hsp_independent_contrasts), which employs a postorder traversal algorithm to predict internal node values and then assigned hidden tip states to their most recent reconstructed ancestor estimated value. Predicted hidden states for continuous data were in the same units as the known host input data. For ordinal data, we used a fixed-rates continuous-time Markov model to estimate transition rates between each ordinal state (castor::hsp_mk_model), which was fit with maximum likelihood. This fitted model was used to calculate marginal likelihoods for each possible state first for internal nodes using the rerooting method proposed by Yang et al.^87^ and then estimated hidden tip states from the most closely reconstructed ancestral node while accounting for the intervening patristic distance via exponentiation of the transition matrix^33 SI,36^. Ordinal data HSP was parameterized under default settings except for as follows: Nstates = 3, rate_model = “ARD”, Ntrials = 1000, optim_max_iterations = 1000, and optim_rel_tol = 1e-10. We selected the number of rates and transition rate model based on the number of states in the ordinal association data and to allow for the expectation that transitions away from use by many SLF life stages is more costly than towards them. Predictions for ordinal HSP then were probabilities for each possible state. For all HSP methods, known host input data remained unchanged in output (treated as known rather than in need of prediction).

SLF host association values used in HSP analyses were natively in different units across studies, making a single determination of SLF association difficult for each species (Table 1). To address this obstacle, we calculated two different composite association metrics, one each for HSP1 and HSP2. To standardize data across studies (*i* = 1, …, *n*) and species (*j* = 1, …, *m*), I rescaled associations, *x_ij_*, to [0,1] as:

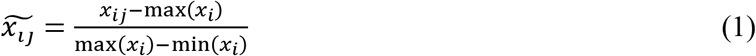

The rescaling step in (1) was conducted for predicted association after HSP but before combining into a composite association metric for HSP1 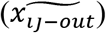 and prior to averaging and HSP for HSP2 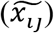. For HSP1, predicted states for our ordinal data required additional steps to obtain a single measure of association from the probabilities for each of the three possible states. Given that both states 2 and 3 represent multiple SLF life stages using a species as a host, we calculated the sum of those two probabilities and used the resulting values as a single measure of association for the ordinal data HSP.

With rescaled associations for each species for each study, we calculated an average association across studies. To do so for HSP1, we computed the product of the HSP output associations 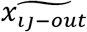 and a contribution weight *w_ij_* = 0.20 that was summed across the five study data sets to obtain a composite SLF association value (analogous to the arithmetic mean) for each candidate host as:

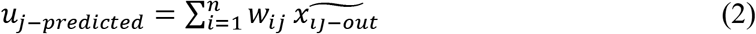

To obtain composite associations for HSP2, I first calculated *u_j_* similarly to *u_j–predicted_* in (2) by replacing 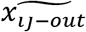 with 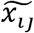 and setting *w_ij_* equal to the reciprocal of the number of studies that include species *j*. Consequently, some species were represented by fewer studies in HSP2 (*w_ij_* was not the same for all species). The resulting averaged known associations were then used to conduct HSP for continuous data to obtain composite predicted associations under HSP2 (also referred to as *u_j–predicted_*).

### Assessing Prediction Accuracy

We first tested for phylogenetic signal for known hosts using known averaged associations from HSP2 (*u_j_*) with Blomberg’s *K*^35^ in the R package phytools^88^ v. 0.7-90. We evaluated significance in *K* via a randomization test with 10000 simulations.

We then assessed the accuracy of the host association predictions with a leave-one-out jackknife procedure^89^ to measure bias for both HSP1 and HSP2 host associations (similarly to^90^). For each jackknife replicate, we fit HSP models by leaving out association data for a single known host from all studies it is found in and estimating predicted associations. For HSP1, we combined the jackknifed HSP predictions with the other studies (ones that predicted the jackknifed species to begin with if that species was not present in all studies) and recalculated *u_j–predicted_*. HSP2 calculations did not require this step, as combining of associations occurs prior to HSP.

Jackknife *u_j–predicted_* were correlated to *u_j–predicted_* for each known host species (Pearson product moment correlation). A strong correlation among the known and jackknifed values for known host species corresponds to high accuracy with low bias prediction, whereas a weak or lack of correlation would suggest that this method has a high bias and does not predict SLF host association well. Bias was also visually inspected by considering the deviations from a 1:1 correspondence between *u_j–predicted_* and jackknife *u_j–predicted_* for known hosts and by comparison of jackknife *u_j–predicted_* values and *u_j–predicted_* for candidate host species. Strong performance of the composite association metric is best described by consistent jackknife *u_j–predicted_* compared to *u_j–predicted_* for known hosts. Additional support for the composite metric is also confirmed by candidate host jackknife distributions that are narrow and in close proximity to their *u_j–predicted_* value.

### Geography of Association

We overlaid the estimated composite SLF host association measures within a geographic context of the contiguous U.S. to better understand how associations may shape SLF’s spread. Current geographic inventories of tree species within the U.S. are available publicly at county level resolution from the USDA PLANTS Database^91^, which we downloaded manually as each raw county-level inventory for each contiguous U.S. state. Each of these raw datasets were cleaned to address erroneous entries, outdated taxonomy, ambiguous taxa, and other database artifacts. Taxonomy was checked against ITIS and updated with the R package taxize similarly to known and candidate hosts datasets to ensure that each species has an accepted Latin name that maps back to all county FIPS codes provided in the inventories. We also included data from USDA NASS Quickstats^82^ to account for regions where cultivated candidate hosts may not have been reported in PLANTS. We also obtained county-level presence data for invasive tree species from EDDMapS^92^ and coordinate records for all species from GBIF^93^, which were aggregated to the county level. All cleaned plant inventories were saved and combined as a single dataset. We intersected this plant inventory with the average of HSP1 and HSP2 estimated host associations and calculated summary statistics for each county to better understand how known and predicted tree hosts are distributed across the newly invaded U.S. range and beyond for SLF.

## Results

### Known Host Association Strengths and Candidate Hosts

Estimation of SLF host associations in the U.S. began with aggregation of known host observations. Given its recent introduction, we included publications both within the U.S. and internationally. As of 2022, we identified 16 candidate SLF association publications, which varied widely in taxonomic scope and data collection methods. After screening and taxonomic updates, we identified five final research articles that assess associations with experimental assays or field surveys for 144 known host species from 43 families (125 and 32 for tree species only, Table 1). Study species coverage varied across the final dataset from 13–127, of which Barringer and Ciafré^54^ was the only dataset with > 50 species measured. We then identified candidate plant hosts by building and refining a working list of tree species in the U.S. Prior to additional analyses, we updated taxonomy for both known and candidate host species datasets, resulting in 569 candidate host species from 79 families and a combined dataset of 743 species from 95 families.

### Host Plant Phylogeny and Association Prediction

Our initial phylogeny of known and candidate host species comprised a tree with 765 terminal tips. We first removed 22 ambiguously coded species (many of which are conspecifics of other species in the host datasets: e.g., *Quercus* sp.), since they lacked known host data. The resulting final phylogeny containing 743 terminal tips, 174 of which are known hosts (Figure 2). Of the tips, 567 species were present within the backbone tree and 176 (26 known host species) were grafted to the tree. To minimize grafting, seven species were recoded to match phylogenetic backbone taxonomy but were returned to their original labeling thereafter.

**Figure 2:**
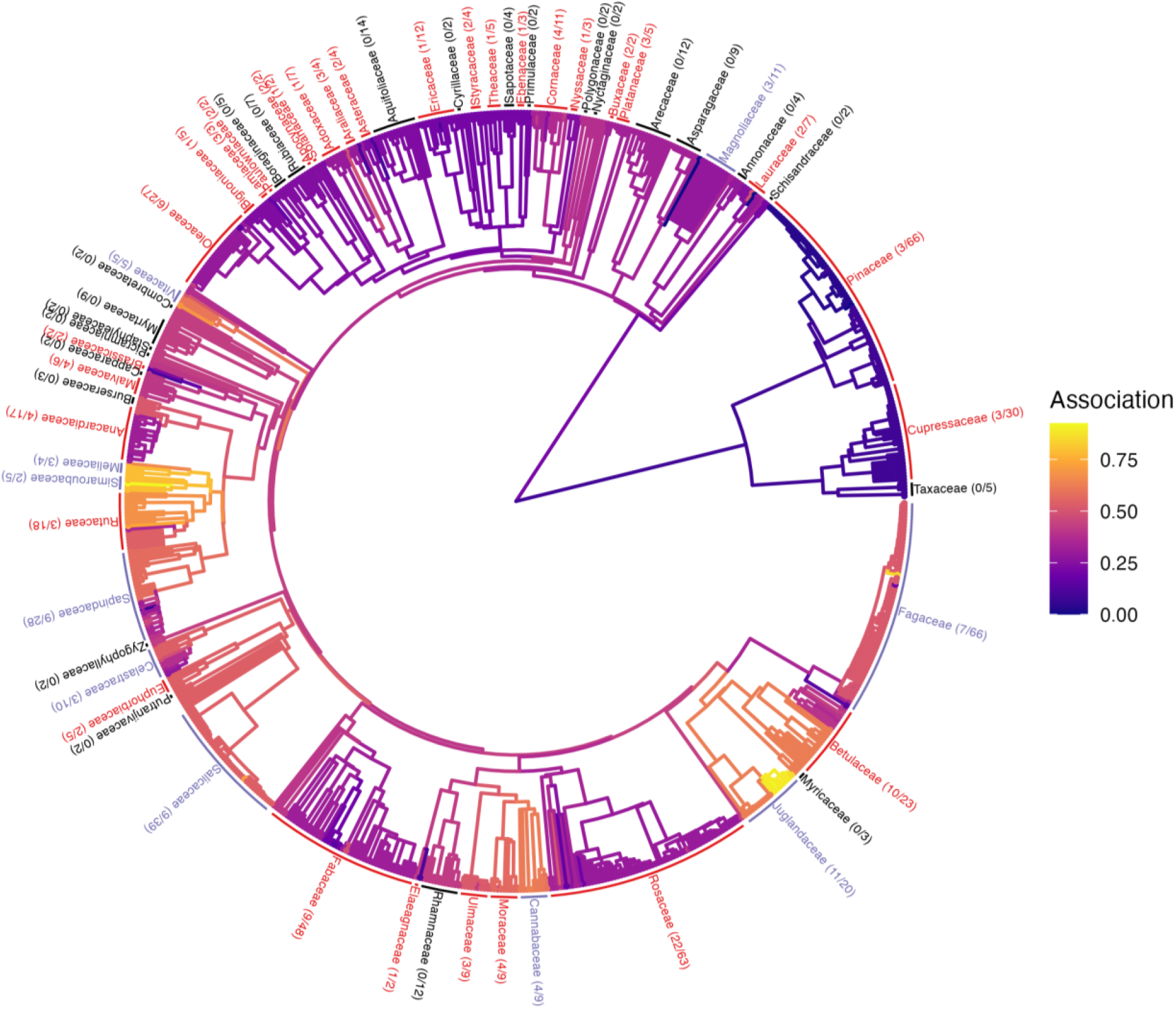
Phylogeny of known and candidate host associations. A phylogeny of plant species that are known or candidate host species for the spotted lanternfly (*Lycorma delicatula*; SLF) shows known and predicted host associations. Host associations are mapped onto branches, with warmer colors indicating stronger association and cooler colors weaker associations. Taxonomic families are labeled that contain at least one known host (red), at least one development host (light purple), and only candidate species (black) at corresponding branch tips and are followed by the number of known host and total species within a family. Some families contain a single species in the phylogeny and are not labeled here.

Estimation of SLF host association was conducted for all species with hidden state prediction (HSP) using two approaches that calculate a composite measure of association. For HSP1, we used all five studies with 144 known host species with data (number of studies: *μ* = 1.4583, σ = 0.8183). Known hosts had a moderate average composite association (*μ* = 0.4276, *σ* = 0.1520) whereas the mean for candidate hosts was slightly lower (*μ* = 0.4161, *σ* = 0.0883). For HSP2, we used the four continuous data studies for 60 known host species (number of studies: *μ* = 1.3833, *σ* = 0.6911), which predicted a similar association relationship but with smaller predicted values for known hosts (*μ* = 0.3620, *σ* = 0.2352) and candidate hosts (*μ* = 0.3579, *σ* = 0.1834).

Although the composite metric was bounded [0,1], these observed associations did not span that full range (Figures 2 and 3). This slightly reduced range for composite associations can likely be attributed to use of an average association that reduces outliers from a study when it is incongruent with all others.

**Figure 3:**
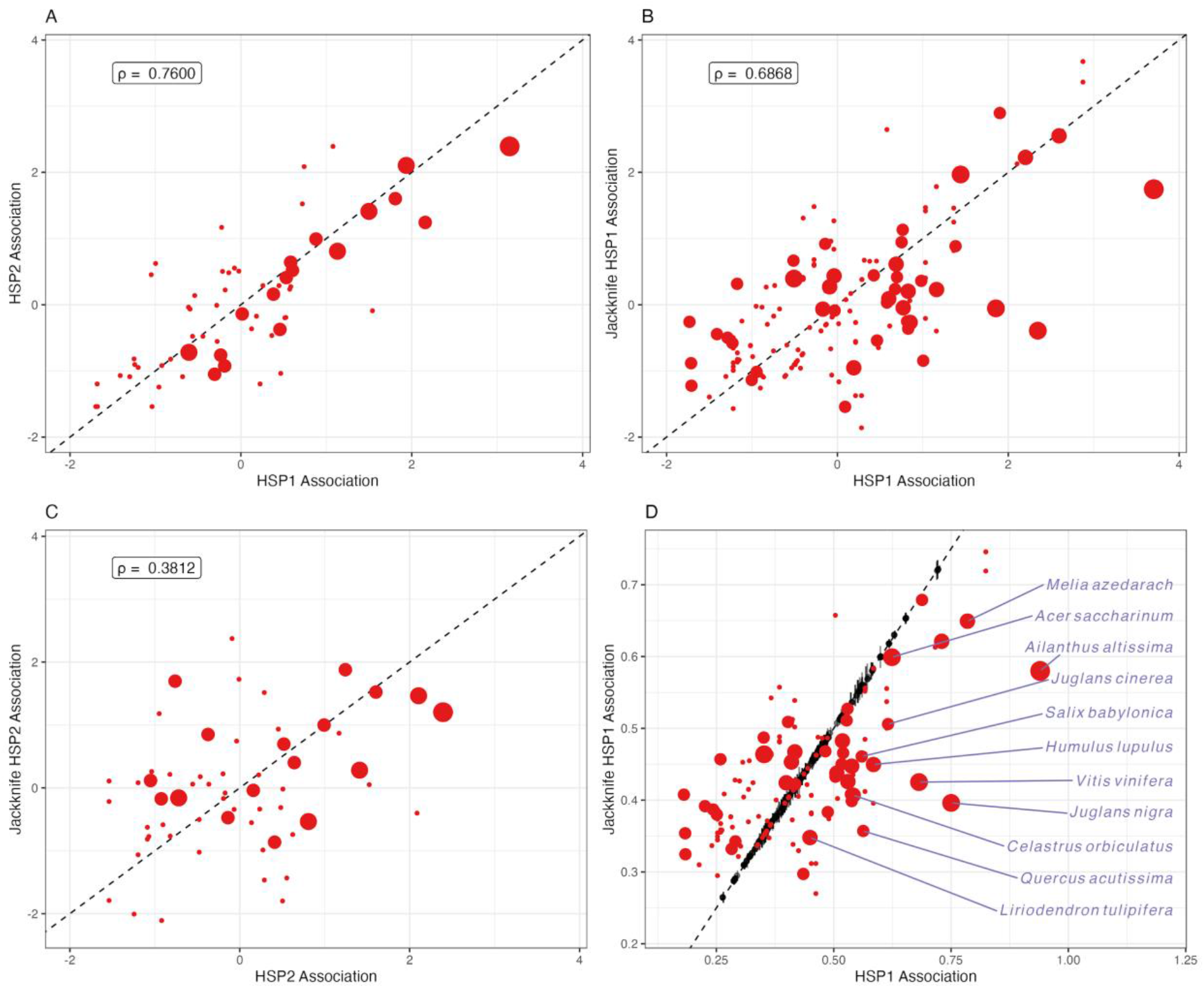
Jackknife and observed host associations. Host associations appear consistent across phylogenetic imputations methods. Observed associations for known hosts (red points) under our phylogenetic imputation models HSP1 and HSP2 were related (**A**). Similarly, known hosts showed a strong relationship between observed values (x-axis) and their jackknife predicted values (y-axis) for HSP1 (**B**) and HSP2 (**C**). For HSP1, both known and candidate hosts had similar values between observed and jackknife replicates (**D**). Candidate hosts were shown as black points for the mean jackknife predicted value (y-axis) and predicted association value (x-axis) with vertical bars indicating the standard deviation for jackknife predicted values. Known host associations along both axes for **A–C** are rescaled and centered, and point size indicates the number of studies that a particular host species is present. Known development host species are labeled for HSP1 (**D**, light purple). The dashed line indicates a 1:1 relationship line drawn through the origin.

### Assessing Prediction Accuracy

To assess the quality of our predictions, we evaluated the phylogenetic signal of host association and prediction bias. Phylogenetic signal was estimated with composite associations for known hosts from HSP2, as they do not include any predictions. We detected weak but significant phylogenetic signal (*K* = 0.2579, *P* = 0.0457). We then evaluated the quality of HSP-predicted SLF host association via a jackknife procedure performed on known hosts. Jackknifed predictions for known host species were obtained via the same methods as other composite association metrics above for HSP1 and HSP2. Known hosts demonstrated a strong correspondence between both methods (Figure 3A, *ρ* = 0.7600, *t* = 8.9056, *DF* = 58, *P* = 1.899e-12). Those found in more studies tended to have lower values for HSP2 compared to HSP1 (Figure 3A). When compared to their jackknife predicted associations, known hosts demonstrated significant relationships for HSP1 (Figure 3B, *ρ* = 0.6868, *t* = 12.393, *DF* = 172, *P* < 2.2e-16) and HSP2 (Figure 3C, *ρ* = 0.3812, *t* = 3.1400, *DF* = 58, *P* = 0.0027), with HSP1 showing a stronger relationship. Jackknife predictions for known hosts indicated that predicted SLF host associations are conservative, as many jackknifed known host predictions were less than their observed counterparts (Figure 3B,C). All but one of the known host trees capable of sustaining SLF to adulthood in laboratory or field conditions were predicted with smaller jackknife associations (Figure 3D purple labels). This observation is consistent with observed composite associations that did not fully extend to rescaled extrema. Regardless of direction, most jackknifed predictions tended to exceed or fall short of observed values by ≤ 0.10, indicating they are similar to observed host associations. Such proximity to observed associations is somewhat surprising, since known host observed associations are not necessarily normally distributed. Rather, numerous known hosts appear to have either very weak or strong association with SLF, which in turn may explain differences between observed and jackknife predicted composite associations (Supplementary Figure 1).

For HSP1, jackknife procedures also produced predictions of candidate hosts without each known host in turn. These jackknife candidate host predictions provided a distribution for devising confidence around predicted associations. Jackknifed predictions of candidate hosts corresponded well with those obtained with the full set of known hosts, with most mean jackknife predictions near those predicted with the full dataset (Figure 3D black points). Furthermore, standard deviation across jackknife replicates was small for candidate hosts, suggesting high confidence in predicted associations (Figure 3D black bars). Additionally, smaller standard deviations indicate that the contribution of each known host to predicted candidate host association does not differ drastically. In other words, no one known host disproportionately determined candidate host composite associations.

### Geography of Association

For application of SLF host association identified here, we considered tree species across the contiguous U.S. To do so, we intersected county-level plant species inventories from the USDA PLANTS database with our combined known and candidate host dataset (filtered for tree family membership for known hosts). Known host species richness per county was greater for those within the current SLF invaded range, including many of the regions that have invested resources into SLF research and management. Altogether, the average number of known tree hosts per county was 18.68 (σ = 14.55) species with the highest number of known host species therein at 70 species and 133 counties reporting no known species, suggesting that the spatial distribution of suitable known hosts is heterogeneous. The addition of candidate hosts yielded a similarly uneven distribution (μ = 55.02, *σ* = 34.61) but with increased species richness for U.S. counties across the eastern midwest and west coast. However, the specific pattern of observed heterogeneity of host species is influenced by underreporting of plant species for counties in IA and MD as well as several counties without any data: Cherokee County, AL, Broomfield County, CO, all parishes of LA, Oglala Lakota County, SD, and Juneau County, WI. Without further data, it may be reasonable to expect that these areas follow neighboring regions and should be assessed when better data are available.

We then used our composite measure of SLF host association averaged across HSP1 and HSP2 to identify regions more accurately in the U.S. with known hosts and predicted hosts with increased likelihood of association. Rather than arbitrarily determine a cutoff for strong association species, we elected to use the lowest observed composite association for a known host species that SLF can reach adulthood on, *Liriodendron tulipifera* (mean association = 0.3893, Figure 3). For this mean threshold, we found that 255 candidate and 70 known host species (64 tree species) species were predicted as strong SLF hosts. This biological threshold reduced the number of suitable host species for each county considerably (Figure 4) for both known hosts (*μ* = 9.772, *σ* = 8.505) and predicted plus known hosts (*μ* = 28.96, *σ* = 20.69). Nevertheless, many of the regions inside and beyond the current invaded range with more naive predicted plus known host species richness demonstrated relatively higher numbers of predicted hosts (Figure 4B). Much of the U.S. midwest, southeast, and west coast contain many species that are predicted to associate with SLF strongly.

**Figure 4:**
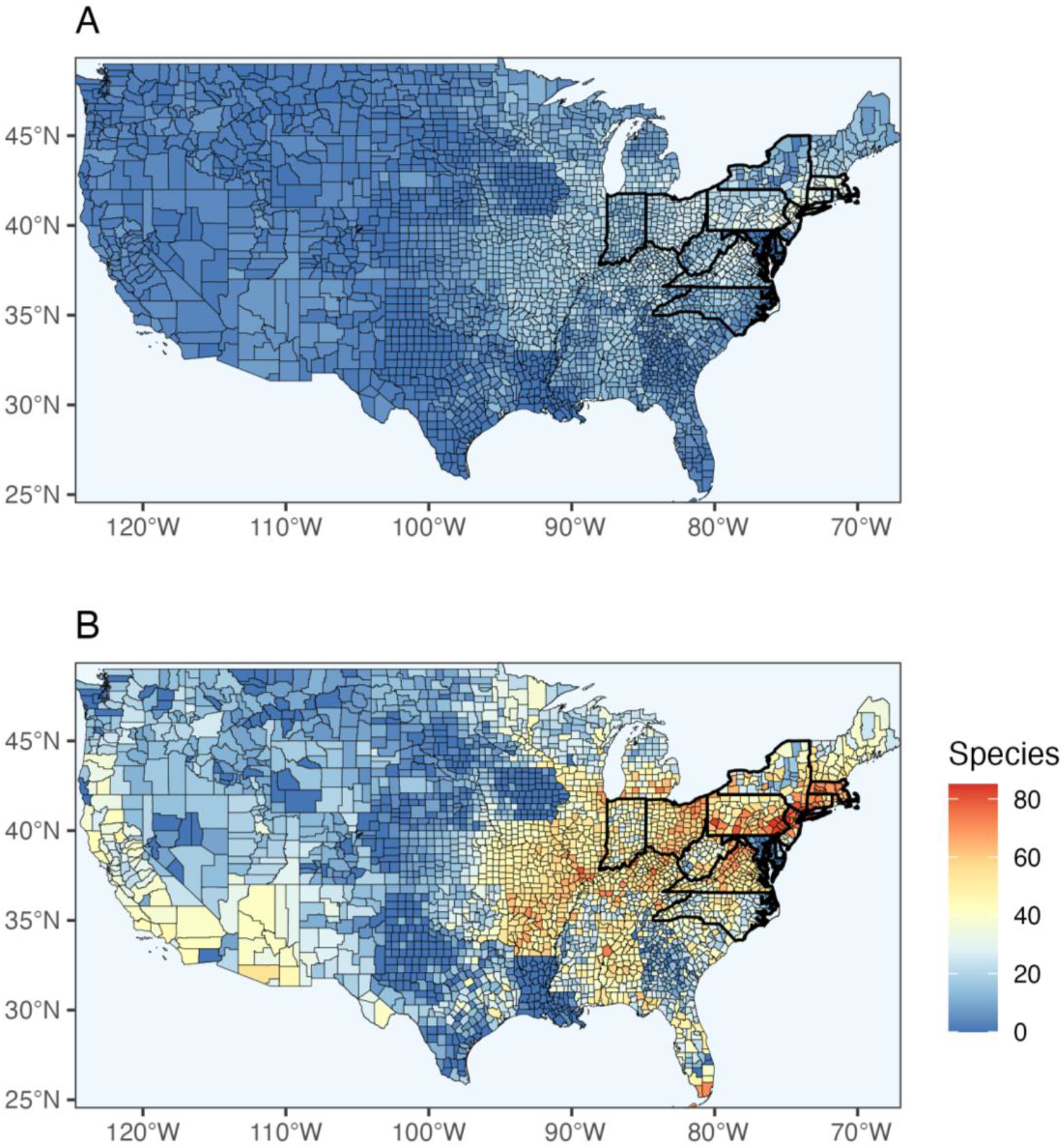
Map of tree species with strong association by county. Species richness of strongly predicted known and candidate host tree species of the spotted lanternfly (*Lycorma delicatula*; SLF) are shown for the contiguous U.S. Biologically informed strongly predicted hosts were set as species with a predicted association greater than or equal to tulip tree, *Liriodendron tulipifera*, the lowest association adult development host species. Strong association known host species (**A**) are mostly restricted to the known SLF distribution (mid-atlantic and northeast, states outlined boldly) and immediately adjacent areas (midwest). When strongly predicted candidate species are added (**B**), many of these same regions are still the most species rich for hosts but with the addition of other regions (southeast and west coast).

## Discussion

SLF is a generalist phytophagous insect pest that boasts an ever-increasing list of hosts, and efforts should be made to better understand the extent and limitations of its host associations to help curtail its rapid spread. When it was first detected in the U.S., SLF was believed to feed on over 70 plant species from 25 families^53^, but since then, the published number has grown to over 100 species from 33 families, including at least 15 species that were not previously known to associate with SLF before research began on the U.S. invaded region^54^. Continued growth in known hosts underscores the importance of continued study of SLF host breadth via field surveys and experimental assays like those used for this study. Although both methods can contribute to our understanding of host associations, we recommend that field surveys be used to direct focus for experimental assays. Host field observations can be documented alongside ongoing federal and state monitoring efforts, making them an inexpensive source of information. For similar reasons, we also encourage citizen science reporting endeavors that capture host association (e.g., ^94^). Documentation of host associations under field conditions is an important early filter, but other methods are necessary to validate and contextualize field data.

In contrast with field observations, comparative association experiments are vital for ground-truthing host associations and determining the factors that drive such preferences. For example, a field observation of a nymph on one tree species may suggest association but not confirm how it is used (e.g., egg laying, feeding, or dispersal) or how association tracks throughout the SLF life cycle^54^. The tradeoff between uncertainty in field observations and limited design capacity of experiments to date (e.g., few host species) motivated our use of both kinds of studies. However, limitations for experiments can be addressed more directly through study design, and so we recommend additional controlled experiments of SLF preference. As resources allow, effective studies should measure fitness-related association consistently (e.g., survivorship rates and per capita offspring production) under natural conditions, prioritize presumptive host diversity, evaluate association throughout SLF’s life cycle, and initially avoid combining hosts in treatments. These design attributes aim to standardize otherwise disparate association units measured (e.g., Table 1) while accounting for key SLF discoveries, such as physiological feeding limits at different life stages^51^, apparent narrowing of host associations as SLF matures^95,63^, and the role of phytochemicals for attraction^96,66^ and toxin sequestration^58^. The ability to account for important covariates in study design is a strength of experiments that makes them ideal for validating the results of other methods like field surveys and predictive modeling when they cannot do so (as is the case here).

Although field surveys and comparative experiments can contribute to our knowledge of insect pests like SLF, they also have limitations that reduce the speed at which we do so. To address emergent pests of concern like SLF, additional approaches like phylogenetic imputation are necessary, even if they must be validated or expanded upon later. If well-constructed phylogenies and association data are available, predictive models that harness them can be inexpensive and fast, especially compared to alternatives like molecular gut analysis^65^ or environmental DNA sampling^97^. In the present study, we demonstrated that SLF host associations show phylogenetic signal and identified potential high-risk hosts without any direct ‘traditional wet lab’ costs. Nevertheless, further elucidation of host association for SLF is hampered due to data availability. For example, we can point to high estimated association for key taxa that contain phytochemicals linked to toxin sequestration in SLF, such as quassinoids in known hosts like *Ailanthus altissima* and *Picrasma quassioides*^98,99^, but we lack information on quassinoid presence in related predicted high association species, such as *Castela emoryi, Leitneria floridana*, and *Simarouba glauca*. Therefore, we can identify that SLF is likely to associate with species in the clade they all belong to, Simaroubaceae. However, why they are suitable for SLF remains unclear, in part because we do not have adequate data for quassinoids across species. The same is true for other clades with many predicted high association species like Juglandaceae and Salicaceae, which are known defensive phytochemical reservoirs^54^. It is clear that better phytochemical data could inform the phylogenetic signal we observed for host association^54^. The same is broadly true for plant trait data^100,101^, which could provide support for alternative drivers of SLF host association like plant structure, sugar content, or sap availability^54,95^. To determine the veracity of these different drivers for SLF and other phytophagous insect pest host associations, gaps in global plant trait data must be filled in.

SLF continues to spread throughout its newly invaded range in the U.S.^49^, and recent spread^50^ corresponds with early exposure to areas that contain species we identify as predicted high suitability hosts, especially in the midwest (Figure 4). Other U.S. regions that are especially vulnerable to SLF because of many known and predicted suitable hosts include regions where SLF is already found (mid-atlantic and northeast) and where it has not yet been found (the west coast and southeast). Regulatory incident reports for several of these not yet invaded regions demonstrate evidence of SLF transportation^49^. However, future investigation of observed geographic patterns in SLF association should include multiple plant inventory sources (e.g., EDDMapS and GBIF) to address insufficient sampling for certain regions here (e.g., all individual parishes in Louisiana and many counties in Iowa). Future robust plant inventories for these regions is necessary. Nevertheless, the currently uninvaded regions above contain strongly predicted host species (members of Anacardiaceae, Betulaceae, Celastraceae, Fagaceae, Juglandaceae, Rutaceae, Salicaceae, and Sapindaceae) in addition to known hosts found there. The U.S. west coast and southeast also contain predicted hosts in Burseraceae and Simaroubaceae, unlike the midwest (known hosts only). Lastly, the southeast (specifically Florida) contains mahogany (*Swietenia mahagoni*, Meliaceae), a predicted host predicted to strongly associate with SLF. We recommend that field monitoring efforts take these regions and taxa into account and that experimental studies prioritize quantifying SLF association strength for representative members of these clades beyond those already sampled for them.

No one approach is a panacea for understanding a generalist insect pest’s host associations, but when used together, associations can become clearer. We have shown here how host phylogenetic relationships and sampled association data can be used together to close gaps in knowledge for a particular pest. Such information can often be obtained inexpensively and more importantly directs management practices and future research using traditional methods for identifying at-risk taxa and regions. Model-based methods are not appropriate for all systems (e.g., not for highly specialized pests) but are useful early in invasions when resources and data are limited or when the biology of a pest is not well understood. Additionally, the research questions about the focal pest must have a scope that phylogenetic relationships can accurately capture and be interpreted cautiously for poor data coverage or in the absence of additional covariates until results can be validated. Despite these limitations, phylogeny-based predictive modeling is a promising tool to inform insect pest surveillance and management.

---

**Supplementary Figure 1:**
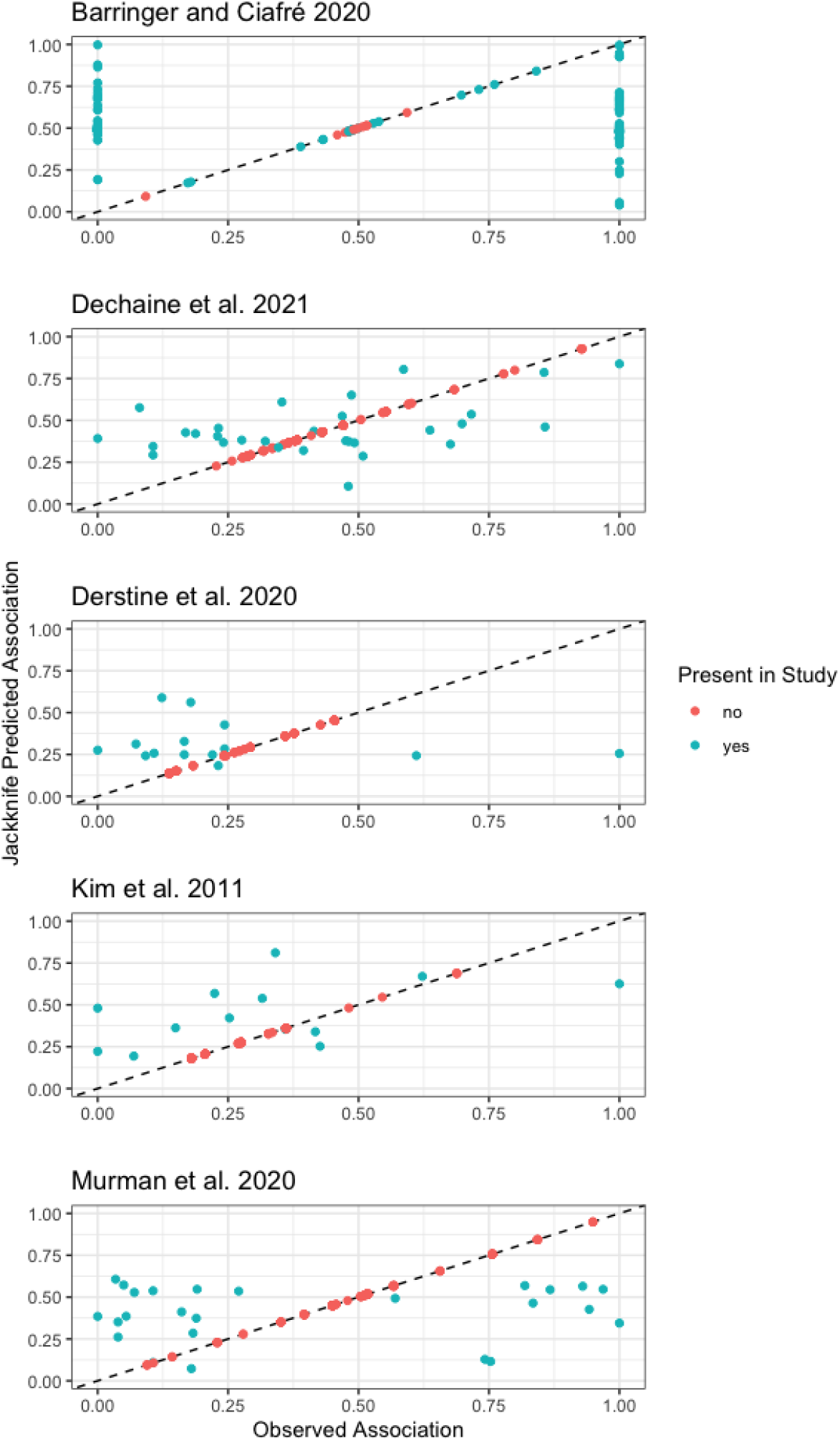
Comparison of observed and jackknife association by study. For individual studies under HSP1 (see main text), the jackknife predicted values are plotted against corresponding observed values. Several studies^54,60,66^ show strong concentrations of observed known host associations that are clustered at distribution extrema but are more diffuse for jackknife predictions.

## Notes

### Competing Interest Statement

The authors have declared no competing interest.

